# Characterization of *Rhodopseudomonas palustris* population dynamics on tobacco phyllosphere and induction of plant resistance to *Tobacco mosaic virus* infection

**DOI:** 10.1101/616110

**Authors:** Pin Su, Deyong Zhang, Zhuo Zhang, Ang Chen, Muhammad Rizwan Hamid, Chenggang Li, Jiao Du, Ju’e Cheng, Xinqiu Tan, Limin Zhen, Zhongying Zhai, Wen Tang, Jin Chen, Xuguo Zhou, Yong Liu

**Affiliations:** Hunan Academy of Agricultural Sciences, Hunan Plant Protection Institute, Changsha 410125, China; Department of Entomology, University of Kentucky, Lexington, KY, 40546, USA; College of Bioscience and Biotechnology, Hunan Agricultural University, Changsha 410128, China

**Keywords:** phyllosphere, colony morphology, biocontrol agent, *Rhodopseudomonas palustris*

## Abstract

Although many biocontrol bacteria can be used to improve plant tolerance to stresses and to promote plant growth, the hostile environmental conditions on plant phyllosphere and the limited knowledge on bacterial colonization on plant phyllosphere minimized the beneficial effects produced by the biocontrol bacteria. *Rhodopseudomonas palustris* strain GJ-22 is known as a phyllosphere biocontrol agent. In this paper we described detailed processes of strain GJ-22 colony establishment at various colonization stages. We have shown that the preferable location sites of bacterial aggregates on leaf phyllosphere are grooves between plant epidermal cells. In this study, we categorized bacterial colonies into four phases. Analyses of expressions of plant defense-related genes showed that, starting from Phase III, bacterial cells in the Type 3 and Type 4 colonies started produce unidentified signals to induce host defense again *Tobacco mosaic virus* infection. To our knowledge, this is the first report focused on the colonization process of a phyllosphere biocontrol agent.

## Introduction

Phyllosphere, also known as aerial part of plant, is the first line plants used to fight against pathogen infections (Berg, 2009; Natacha et al., 2014). Phyllosphere also provides an ideal habitat for bacterial species that are beneficial to plant growth and stress tolerance (Duke et al., 2017; Kembel et al., 2014; Vorholt, 2012). Successful applications of microbes with biocontrol activities to plant phyllosphere are crucial for effective crop disease managements (Kim et al., 2011; Lambais et al., 2014). The most commonly used microbes with biocontrol activities are plant growth-promoting rhizobacteria (PGPR), and the most well-known PGPRs are those currently being used to control crop soil-borne diseases (Bulgarelli et al., 2013; Hassani et al., 2018). Introduction of PGPRs to plant phyllosphere is not easy, due mainly to the hostile environments, including limited nutrients, limited water supply, UV radiation, and toxic substances secreted by plants or produced by other microbes (Lindow & Brandl, 2003; Müller et al., 2016). These hostile conditions can dramatically affect PGPR survival and population density in field. Earlier studies had indicated that inhibition of beneficial microbe growth could lead to much lower microbial population density on leaf phyllosphere and thus significantly reduce the beneficial effects caused by PGPR on plant growth and/or defense response against stresses (Bonaterra et al., 2007). Consequently, we reason that increase of microbial population densities on leaf phyllosphere should enhance the beneficial effects caused by PGPR on plant. To our knowledge, the current knowledge on bacterial colonization on leaf phyllosphere is mostly from plant pathogenic bacteria such as *Pseudomonas syringae* (Haefele & Lindow, 1987) and *Erwinia* (*Pantoea*) spp. (Okinaka et al., 2002). The current knowledge on PGPR colonization on leaf phyllosphere is very limited.

*Rhodopseudononas palustris* is a purple non-sulfur photosynthetic bacteria in Class *Alpha-proteobacteria*, Phylum *Proteobacteria*. Strains of *R. palustris* are often known as free-living bacteria in soils or in aquatic sediments (Larimer et al., 2004). Strains of this bacterium had drawn enormous interests from researchers who are interested in *R. palustris* metabolic abilities, including nitrogen fixation, hydrogen gas production, and degradation of aromatic compounds and chlorinated pollutants (Sasaki et al., 2005). *R. palustris* is now widely used to treat agricultural and industrial wastes or to produce bioenergy (Idi et al., 2015). Its potentials in organic farming are new topics of many research laboratories (Fawzy et al., 2012; Wong et al., 2014; Wu et al., 2013). In our previous report, we have shown that *R. palustris* GJ-22 strain can induce a systemic resistance (referred to induced systemic resistance [ISR] thereafter) to *Tobacco mosaic virus* (TMV) infection in tobacco and promote plant growth through productions of two phytohormones (i.e., indole-3-acetic acid and 5-aminolevulinic acid) (Su et al., 2017). Unlike many other PGPR strains that colonize plant rhizosphere, *R. palustris* GJ-22 can colonize plant phyllosphere and establish different sized colonies to exert beneficial effects to plant. After being introduced to tobacco phyllosphere, the population density of *R. palustris* GJ-22 declined quickly and then recovered to a form large sized colonies. This phenomenon suggests that *R. palustris* GJ-22 utilizes some unidentified strategies to overcome biotic and abiotic stresses during its on plant phyllosphere. Despite the reported beneficial effects of *R. palustris* strains and their ability to colonize phyllosphere, an understanding of colonization pattern in the phyllosphere is a critical prerequisite when deployed into field as an biocontrol agent. Strains of *R. palustris* are often referred to as free-living bacteria in soils or in aquatic sediments. Current knowledge on *R. palustris* lifestyle are largely from the soil or aquatic *R. palustris* strains. There is little known about how *R. palustris* adapt to phyllosphere conditions.

In this study, we investigated the formation of bacterial colony on tobacco leaves inoculated with strain GJ-22-eGFP, the wild type strain GJ-22 expressing a free eGFP protein, using both Confocal Laser Scanning Microscopy (LSCM) and Scanning Electron Microscopy (SEM). We found that *R. palustris* strain GJ-22 could form Type 3 colony by 72 h post bacterial inoculation and Type 4 colony by 96 h post bacterial inoculation on leaf phyllosphere. *R. palustris* GJ-22 cells inside the Type 3 and 4 colonies were able to induce ISR in tobacco to *Tobacco mosaic virus* (TMV) infection. We consider that the findings presented here provide useful information on the establishment of *R. palustris* colonies on leaf phyllosphere and information needed for the development of new strategies using this biocontrol agent.

## Materials and methods

### Bacterial strains, cultural conditions and plant inoculum

*R. palustris* strain GJ-22 was from a previously reported source (Su et al., 2017) and routinely cultivated aerobically at 30 °C in a photosynthetic medium (PM) under a 6,500 lux lamination. The solid PM medium contains 0.1 g (NH_4_)_2_SO_4_, 0.02 g MgSO_4_, 0.5 g Na_2_CO_3_, 0.05 g K_2_HPO_4_, 0.02 g NaCl, 0.2 g Casamino Acids, 0.15 g yeast extract and 18g agarose in one litre distilled water, pH 6.5-7.0. For liquid PM medium, agarose was omitted from the above formula. *Escherichia coli* DH5α was aerobically cultured in the Luria-Bertani (LB) broth at 30 °C and used for plasmid propagations.

The transformation of GJ-22 wild type with plasmid pBBR1MCS-2-pAMP-EGFP (Beijing TsingKe Biotch Co., Ltd., Beijing, China) was conducted through the following procedure. Cells were pelleted from liquid culture (OD660 = 0.4), rinsed three times with an ice-cold sterile water before resuspended in 10 % glycerol solution, and resuspended cells was immediately stored at −80 °C for later use. Electro-competent cell suspension (50 μL) was thaw on ice for 20 mins, mixed with 5 ng plasmid DNA in a 2 mm Gene Pulser cuvette containing 80 μL ice-cold sterile water, incubated on ice for 10 mins for incubation followed by electroporation using Eppendorf Eporator as described (Eppendorf North America, Hauppauge, NY, USA). The transformed cell suspension was transferred to 10 mL liquid medium and cultured for 20 h at 30 °C with 6,500 lux illumination inside a light incubator. The positive transformants were selected on agar medium with 50 µg·mL^−1^ kanamycin under the same culture condition described above. Stability of the mutants was analyzed by cultivating them individually in the liquid medium, with and without kanamycin, for 20 generations. For EGFP labeled strain GJ-22-EGFP, the fluorescence shown by each generation was examined with a laser scanning confocal microscope (LSCM) (Nikon C2 plus, Nikon, Japan). The excitation wavelength was set at 488 nm and the emission wavelength was set at 522/35 nm.

To prepare fresh bacterial culture, small amount of original cell stock culture (OD_660_ = 0.4) was added into a fresh liquid medium at the ratio of 1:20 (stock bacterium: medium, v/v). Before bacterial inoculation, GJ-22 or GJ-22-EGFP culture was pelleted, resuspended in a 100 mM potassium phosphate butter (PPB), pH 7.2, and adjusted to 6 × 10^7^ CFU mL^−1^ immediately prior to inoculation to plant.

### Plant growth and inoculation

Seeds of *Nicotiana. benthamiana* were placed in a 70 % ethanol solution for 30 min at room temperature (RT). After removing 70 % ethanol by low speed centrifugation, the seeds were treated with 100 % ethanol for 15 min. The seeds were dried on filter papers for 1 h and geminated on wet filter papers at 25 °C. The germinated seeds were individually transferred to pots (10cm × 10cm × 15cm) containing pH-balanced peat moss (Klasmann-Deilmann GmbH, Geeste, Germany) inside a growth chamber set at 28 °C, a 16/8 h (light/dark) photoperiod, and 85 % relative humidity. For stress assays, the growth chamber was set at 37 °C, a 16/8 h (light/dark) photoperiod, and 40 % relative humidity. Bacterial inoculation was conducted by spraying 14-day old *N. benthamiana* seedlings till the seedling became soaking wet. The inoculated seedlings were grown inside a growth chamber the conditions described above.

### Bacterial sampling and phyllosphere population density determination

Bacterial cells were collected at various hours post plant inoculation (hpi) with the following method. The third and fourth leaves (down from the top) were harvested from individual assayed plants and pooled. Ten gram tissues were sampled from each harvested leaf sample and submerged in 200 mL PPB inside a 500 mL conical flask. After 20 min, the samples were sonicated at 47 kHz for 10 mins and with 150 rpm shaking. Bacterial cells in each PPB solution were pelleted through 10 min centrifugation at 10,000 *g*. This procedure was repeated three times and the bacteria containing PPB solutions from each leaf sample were pooled. After centrifugation, the pelleted cells was resuspended in 2 mL PPB with 20 % Percoll followed by 10 min centrifugation at 12,000 *g* to remove the remaining plant debris.

Bacterial population density in each sample was determined by plating the serially diluted bacteria containing solutions on PM plates. The inoculated plates were then incubated for 6 days at 25 °C and the colony-forming units (UFC) on each plate was counted and later converted to UFC g^−1^ FW (bacterial population size).

### Assay of ISR onset in *R. palustris* strain GJ-22 inoculated *N. Benthamiana*

PPB solution containing strain GJ-22 was sprayed onto the fifth and sixth leaf of *N. Benthamiana* seedlings. Seedlings sprayed with equal amount of PPB only were used as controls. The sprayed leaves were detached from the plants at 12, 48, 72, and 96 hpi. At 12 h post detachment, the third and fourth leaf of each seedling were mechanically inoculated with purified TMV (40 ng per leaf). The TMV-inoculated leaves were harvested at 6 days post TMV inoculation and analysed for TMV accumulation by Enzyme-linked immunosorbent assay (ELISA). To determine the expressions of defence-related genes, the TMV-inoculated leaves were harvested at 18 hours post TMV inoculation and analysed for the expressions of defence-related protein genes *NbPR1a* and *NbPR3*, through semi-quantitative reverse transcription polymerase chain reaction (qRT-PCR) as described previously (Su et al., 2017).

### Confocal Laser Scanning Microscopy (LSCM) and Scanning Electron Microscopy (SEM)

For LSCM analysis, leaf tissues were collected and cut into 2 × 2 mm pieces. Bacterial cells on the surface of each leaf sample was examined under a Nikon epifluorescence microscope confocal system (Nikon TI-E+C2, Japan). The excitation wavelength was set at 488 nm and the emission wavelength was set at 522/35 nm. The SEM analysis was conducted as described by Poonguzhali et al. (Beattie & Lindow, 1999; Poonguzhali et al., 2008) with minor modifications. Leaf tissues were collected and cut into 0.5 cm^2^ pieces. The tissues were transferred into a 0.1 M phosphate buffer (PB) containing 2.5 % glutaraldehyde and incubated for 40 min at RT followed by 1 h incubation in a 1 % osmium tetroxide solution at RT. After three rinses in 0.1 M PB, the fixed tissues were dehydrated in 30, 40, 50, 60, 70, 80, 90 and then 100 % ethanol solution. After dehydration, the tissues were freeze-dried and coated with gold-palladium using a Pelco 3 sputter coater (PELCO, USA) and examined under a scanning electron microscope (Nova NanoSEM230, USA).

### Statistical analysis

All the experiments performed in this study were done in three parallels. Results shown in each figure or table were from a single experiment. The results were presented as the mean ± standard deviation (SD) from three or four biological replicates. Statistical differences between the treatments were determined using the One-way ANOVA followed by the Tukey’s test using the SPSS Statistics 17.0 software (IBM Corp., New York, USA).

## Results

### Population dynamic and strain GJ-22 colony morphology on *N. Benthamiana* phyllosphere

Analysis of population dynamic of *R. palustris* strain GJ-22 on *N. Benthamiana* phyllosphere demonstrated that strain GJ-22 took 96 h to reach full population size (Fig. 1). Within the first 24 h, the population size dropped from 5.3 × 10^7^ CFU mL^−1^ to 4.8 × 10^3^ CFU mL^−1^. This population size maintained for at least 24 h and then increased from 68 hpi (3.14 × 10^4^ CFU mL^−1^) to 84 hpi (4.63 × 10^6^ CFU mL^−1^). This steady increase continued in the following 12 h and reached to 9.7 × 10^6^ CFU mL^−1^ by the 96 hpi. The population dynamic can be summarized as a dramatic decline during early growth stage followed by a static state, a slow increase, and then a rapid increase (Fig. 1). Similar dynamic patterns were observed in all parallel experiments.

**Figure 1.**
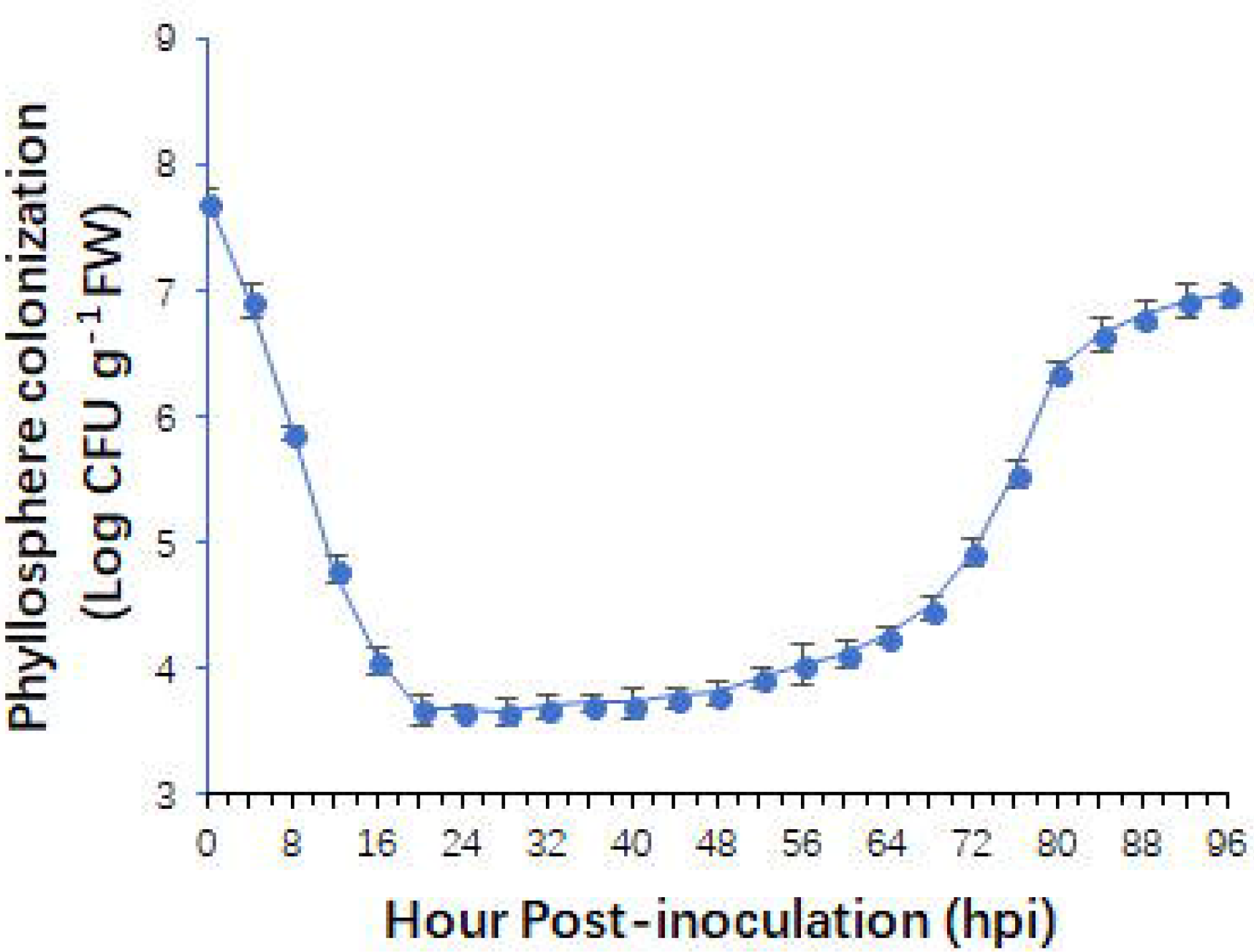
Population dynamic of *R. palustris* strain GJ-22 on *N. Benthamiana* phyllosphere. *R. palustris* strain GJ-22 was inoculated onto *N. Benthamiana* leaves. Population densities of strain GJ-22 on *N. Benthamiana* phyllosphere were examined at 4 h intervals from 0 ~ 96 hpi. The plant growth conditions were set at 28 °C, 16/8 h (light/dark) photoperiod, and 85 % relative humidity. The data were presented as the means from four biological repeats. Error bars represented the standard deviation.

Analysis of green fluorescent signal produced by strain GJ-22-eGFP through LSCM or analysis of wild type GJ-22 through SEM showed that the bacterial cells on phyllosphere formed four types of colonies during colony establishment (Fig. 2). At the initial colonization state (0 to 20 hpi), bacterial cells were mostly exited as single cells randomly distributed on leaf surfaces. No typical bacterial cell arrangements and locations on phyllosphere were observed. Representative images captured by LSCM or SEM at 12 hpi are shown as Type 1 in Figure 2. Single bacterial cells on leaf surfaces died over time, leading to a decrease of green fluorescent signal under LSCM. The death of bacterial cells was likely the cause of rapid decrease population size during this period. The cells lodged at the junctions between plant epidermal cells survived and multiplied afterward. Between 20 and 48 hpi, the multiplied cells began to form small clusters at the junctions. Representative images captured at 48 hpi are shown as Type 2 in Figure 2. Small clustered bacterial cells continued to grow and form aggregates during the next 24 h. Representative images captured at 72 hpi are presented as Type 3 in Figure 2. The cell aggregates enlarged rapidly to form large aggregates. By 96 hpi, the bacterium existed mostly as large aggregates. Representative images captured at 96 hpi are presented as Type 4 in Figure 2. At this time point, bacterial colonies were no longer confined at the junctions between plant epidermal cells but expanded to the surrounding surfaces. In summary, four different types of bacterial colonies: Type 1, scattered single cells; Type 2, small cell clusters; Type 3, small cell aggregates and Type 4, large cell aggregates, were observed in this study.

**Figure 2.**
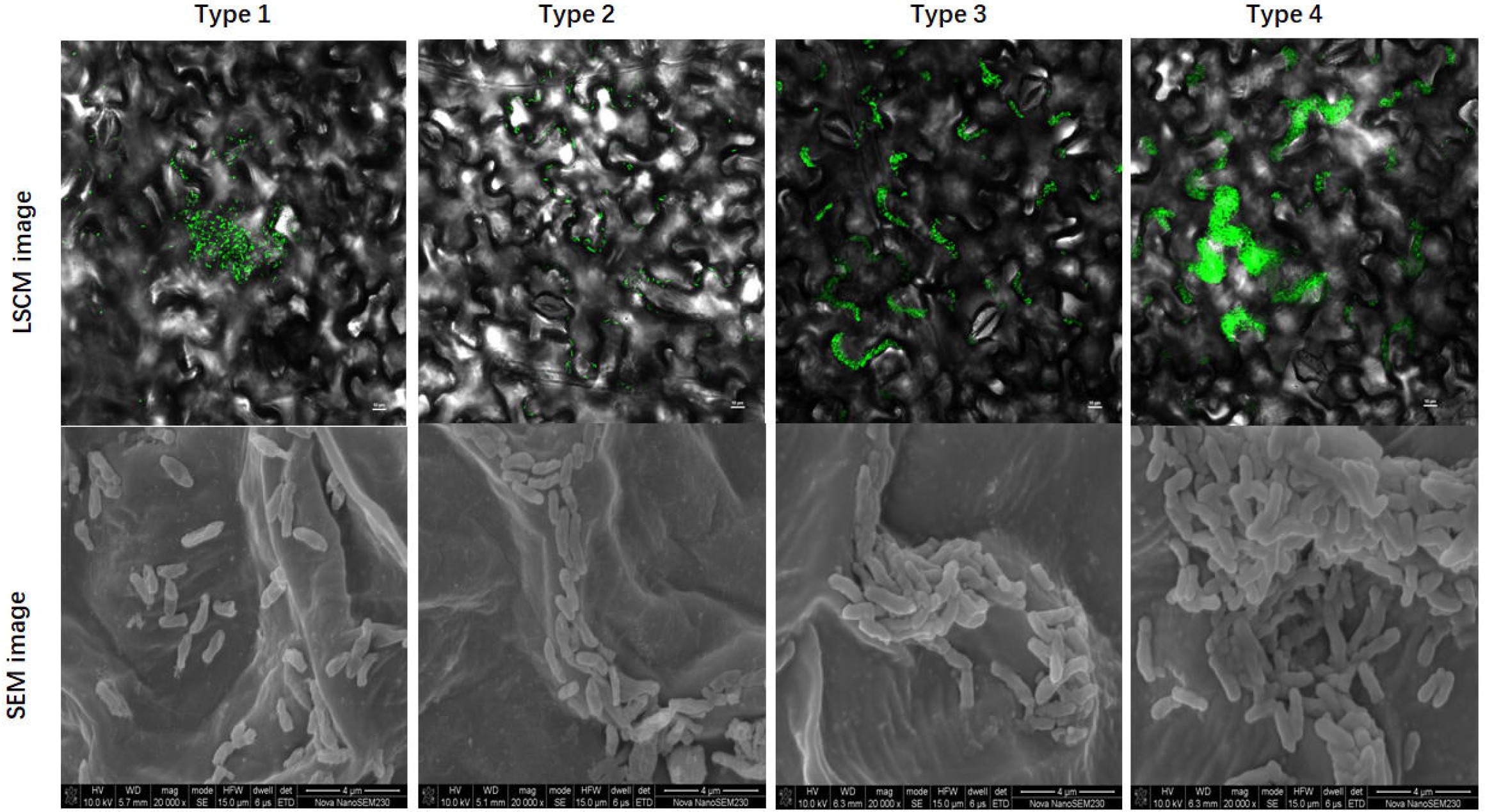
Distributions and colony morphology of *R. palustris* strain GJ-22 on *N. Benthamiana* phyllosphere. The GFP fluorescent produced by GJ-22-eGFP, a wild type GJ-22 harboring an eGFP gene, was sprayed onto *N. Benthamiana* leaves. Images of bacterial cells were captured by LSCM (upper panel) or SEM (lower panel) at *12* hpi (Type 1), 48 hpi (Type 2), 72 hpi (Type 3) and 96 hpi (Type 4), respectively. *Bars in upper panel = 10 μm, bars in lower panel = 4 μm.*

To further illustrate colony dynamics from Type 1 to Type 4, we calculated the percentage of leaf samples showing each colony type (Fig. 3). In the early stage of colonization (0 ~ 20 hpi), over 70 % leaf sample had the Type 1 colonies. Type 2 colonies started to appear at 16 hpi and by 48 hpi, 72 % leaf sample had Type 2 colonies while the Type 1 colony was observed in only about 8 % leaf samples. The number of Type 2 colony started to decrease afterwards and the Type 3 colony started to emerge. By 72 hpi, about 60 % leaf sample contained Type 3 colonies while only 6 % leaf sample contained Type 2 colonies. By 96 hpi, about 60 % leaf sample contained Type 4 colonies while only 4 % leaf sample contained Type 3 colonies (Fig. 3).

**Figure 3.**
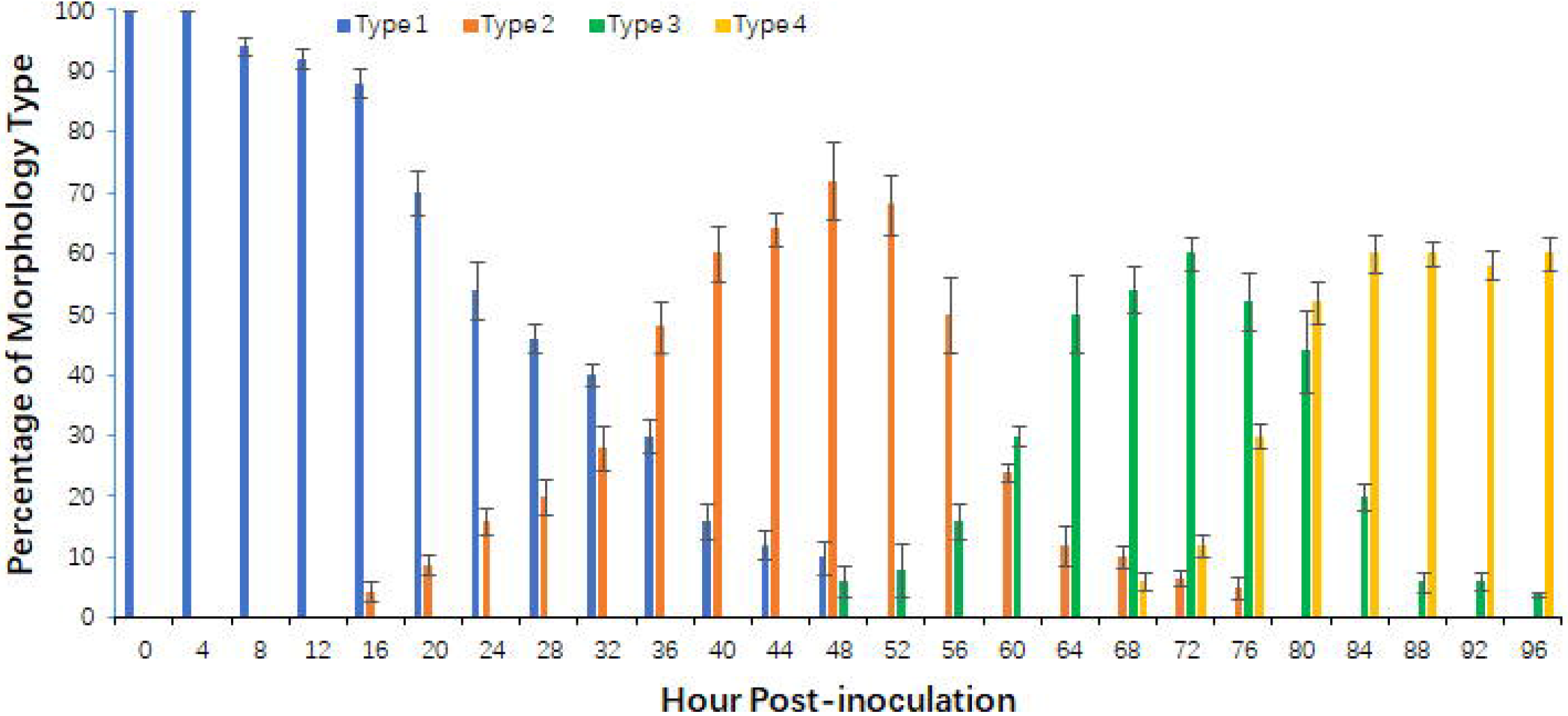
Morphological changes from Type 1 colony to Type 4 colony during the colony establishment. GJ-22-eGFP cells were sprayed onto *N. Benthamiana* leaves. The inoculated leaves were collected at 4 h intervals from 0 ~ 96 hpi. Fifteen leaf discs (2 × 2 mm) were randomly sampled from leaves of each plant (three plants per treatment). The collected leaf samples were examined for bacterial colony morphology by LSCM. The number of samples carrying a specific colony type was recorded and used to calculate the percentage of samples with a specific colony type. The results are presented as the mean percentage ± standard deviation.

### Formation of aggregates enhanced strain GJ-22 tolerance to drought and heat stresses

To investigate whether formation of bacterial aggregate could enhance GJ-22 tolerance to stresses, we treated bacterial cells in different colony types with drought and heat stresses. *N. Benthamiana* seedlings were inoculated with GJ-22-eGFP cells and then grown under the normal growth condition (Fig. 4A). After 12 h (leaves contained mostly Type 1 colonies), 48 h (leaves contained Type 2 colonies), 72 h (leaves contained mostly Type 3 colonies) and 96 h (leaves contained mostly Type 4 colonies), the inoculated plants (4 plant per time point) were transferred to stressed condition (Relative humidity = 40 %, Temperature = 39 °C). The stressed plants were sampled and examined by LSCM. Results showed that the Type 3 and Type 4 colonies maintained their morphology after 1 or 3 day stress treatment (Fig. 4A). The Type 1 and Type 2 colonies died rapidly after the treatment. Analysis of population densities after the stress treatment showed that cells in the Type 3 and Type 4 colonies were more tolerant to drought and heat stresses than the cells in the Type 1 and Type 2 colonies, based on the drastic cell density drop in Type 1 and Type 2 colony and the cell density in Type 3 and Type 4 colony maintained at similar level after stress treatment (Fig. 4B).

**Figure 4.**
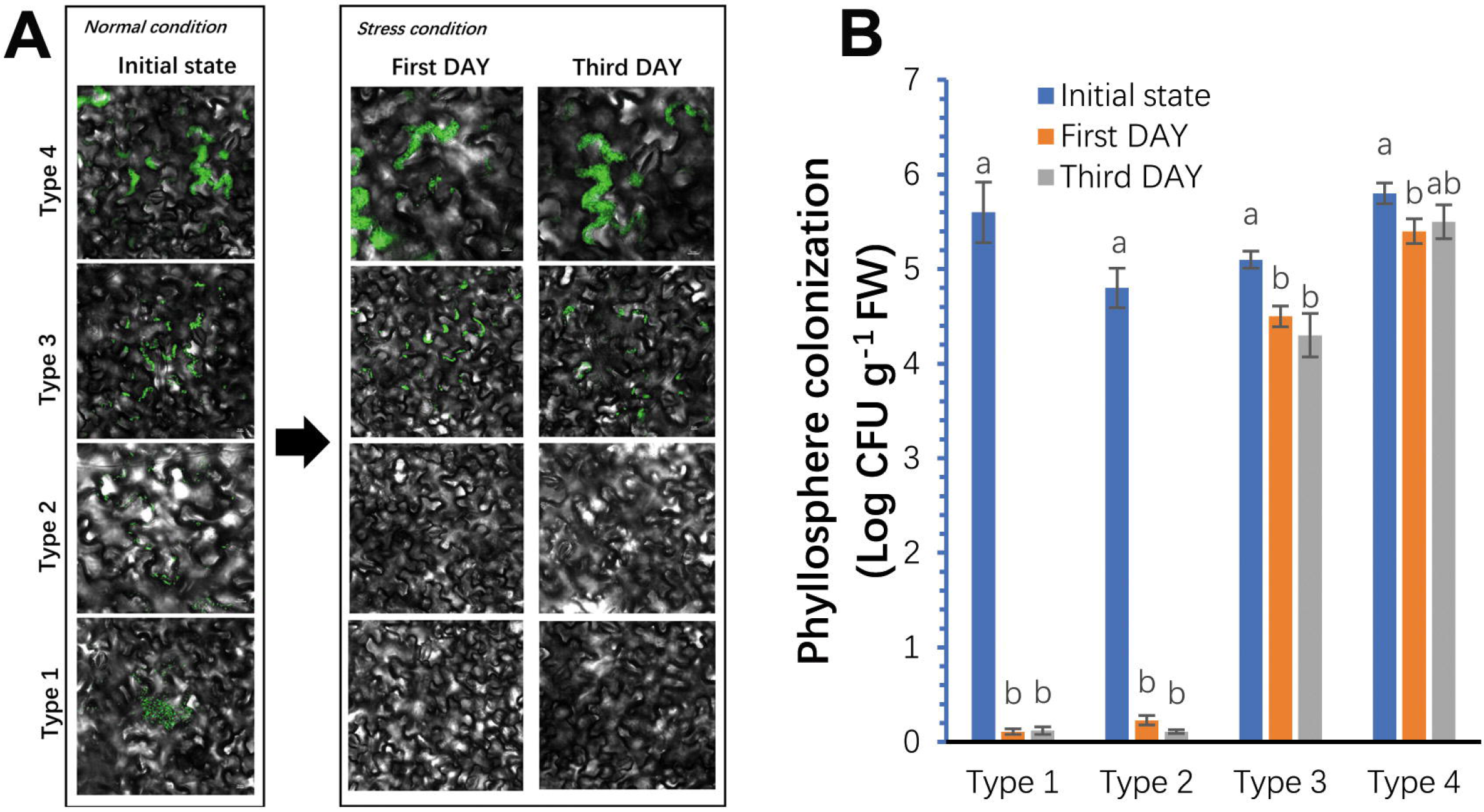
Stress tolerance of strain GJ-22 in different types of colonies on *N. Benthamiana* phyllosphere. (**A**) LSCM images of strain GJ-22-eGFP on *N. benthamiana* phyllosphere under different growth conditions. Bacterial cells on the plants grown under the normal conditions were labeled as initial stage. Representative images of GJ-22-eGFP on plant phyllosphere at 12 (Type 1), 48 (Type 2), 72 (Type 3), and 96 (Type 4) hpi are shown. After being grown under the normal condition, the plants were grown under the stress conditions for three days before LSCM. Images were taken from these plants at one or three days post stress treatment. (B) Population density of strain GJ-22 on *N. benthamiana* phyllosphere under different growth conditions. Population density was determined at the initial state, and first and third day post stress treatment. The data presented were the means ± standard deviations from four biological replicates per treatment. Different small letters above the bars indicate the statistical difference between treatments within the same group by the Fisher’s LSD (*p* = 0.05).

### *R. palustris* strain GJ-22 primed ISR in plants after aggregate formation on phyllosphere

Timing of ISR onset during strain GJ-22 colonization process was investigated. The GJ-22-eGFP inoculated seedlings were grown under the normal condition for 12, 48, 72 or 96 h to produce Type1, Type 2, Type 3 and Type 4 colonies (Fig. 4A). After removal of the inoculated leaves, the assayed plants were inoculated with purified TMV (Fig. 5A). Accumulation of TMV in the inoculated plants was determine through ELISA at 6 dpi, to evaluate the ISR against TMV. In the plants had Type 3 or Type 4 GJ-22 colonies, the accumulation levels of TMV were 52 and 67.4 % lower than that in the plants without *R. palustris* strain GJ-22 inoculation (Mock) (Fig. 5B). For plants had Type 1 and Type 2 colonies, the accumulation levels of TMV were similar to that in the Mock control plants. We then analyzed the expressions of *NbPR1a* and *NbPR3* in plants at 18 h post TMV inoculation to validate the onset of ISR (Fig. 5C). The plants had Type 3 or Type 4 colonies showed quicker upregulations of *NbPR1a* and *NbPR3* expressions compared with the mock plants or the plants had Type 1 or Type 2 colonies, suggesting that strain GJ-22 had the ability to prime ISR in plants after formation of Type 3 and Type 4 colonies.

**Figure 5.**
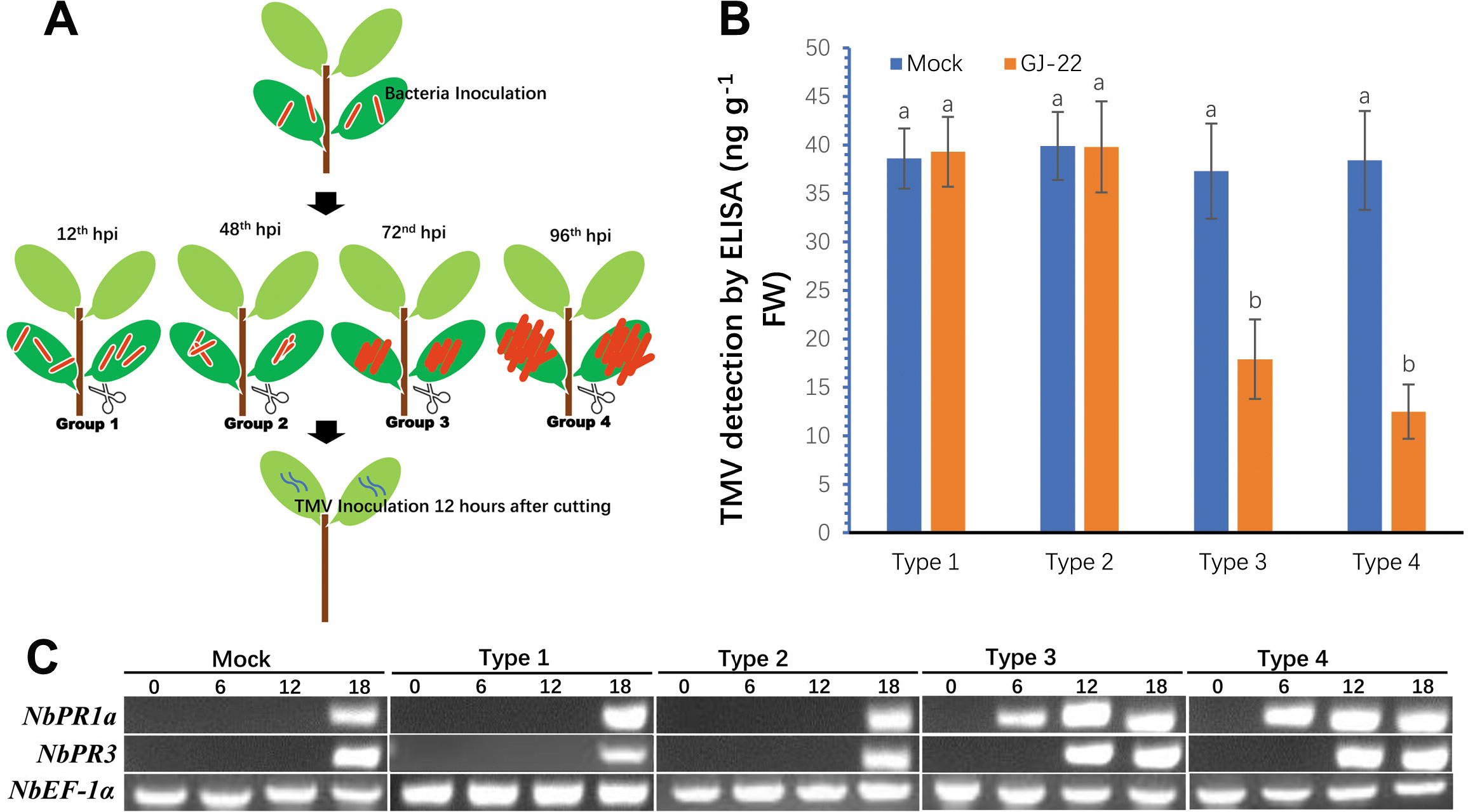
Priming ISR onset by *R. palustris* strain GJ-22 in *N. benthamiana* plants. (**A**) A schematic diagram showing experimental design. The fifth and sixth leaves of *N. benthamiana* plants were sprayed with bacterial strain GJ-22 suspension. After detachment of the sprayed leaves at 12, 48, 72 or 96 hpi, the plants were inoculated with TMV. The TMV inoculated leaves were harvested at 6 dpi. (**B**) Accumulation of TMV in the inoculated leaves was determined by ELISA. The results were presented as the means ± standard deviations from four biological replicates per treatment. Different small letters above the bars indicate statistical differences between treatments in each group by the Fisher’s LSD (*p* = 0.05). (**C**): Induction of NbPR1a and NbPR3 expressions in the TMV-inoculated leaves. The TMV-inoculated leaves were harvested at various hpi and analyzed by semi-quantitative RT-PCR followed by electrophoresis. The expression of NbEF-1α was used as an internal control. Similar results were obtained in three parallel experiments.

## Discussion

In this report we have demonstrated that after attachment to tobacco phyllosphere, *R. palustris* strain GJ-22 underwent a gradual change of colony morphology, starting from Type 1 colony to Type 4 colony. The change from Type 1 to Type 2 colony took about 36 h, and change from Type 2 to Type 3 colony or Type 3 to Type 4 to about 24 h. In addition, colony morphology change accompanied with specific bacterial population densities. For example, change from Type 1 to Type 2 colony occurred during the static state of bacterial growth while the change from Type 2 to Type 3 colony occurred during the bacterial slow growth stage. When strain GJ-22 started to grow rapidly on tobacco leaf phyllosphere, Type 4 colony became to appear. Based on these findings, we characterized strain GJ-22 colonization on leaf phyllosphere into four different phases (Fig. 6). In the phase I stage, individual strain GJ-22 cells are randomly distributed across the leaf surface. These cells die quickly, due mainly to environmental stresses, leading to quick reduction of cell population and colony size. In phase II stage, strain GJ-22 cells started to form small clusters located at the junctions between plant epidermal cells. At this stage, bacterial cell population density remained steady and at the lowest level. In the phase III stage, strain GJ-22 cell clusters started to expend and formed small aggregates at the junctions between plant epidermal cells. Bacterial cells in small aggregates or small colonies started to produce unidentified substances that could enhance plant tolerance to biotic and abiotic stresses. The size of bacterial aggregate or colony continued to expend slowly in this phase. In the phase IV stage, the bacterium formed large aggregates or colonies on leaf phyllosphere and the size of bacterial colony continued to expend due to rapid cell multiplications. By the end of this phase, bacterial aggregates started to emerge to form very large aggregates or colonies, companied with an significant increase of bacterial population density.

**Figure 6.**
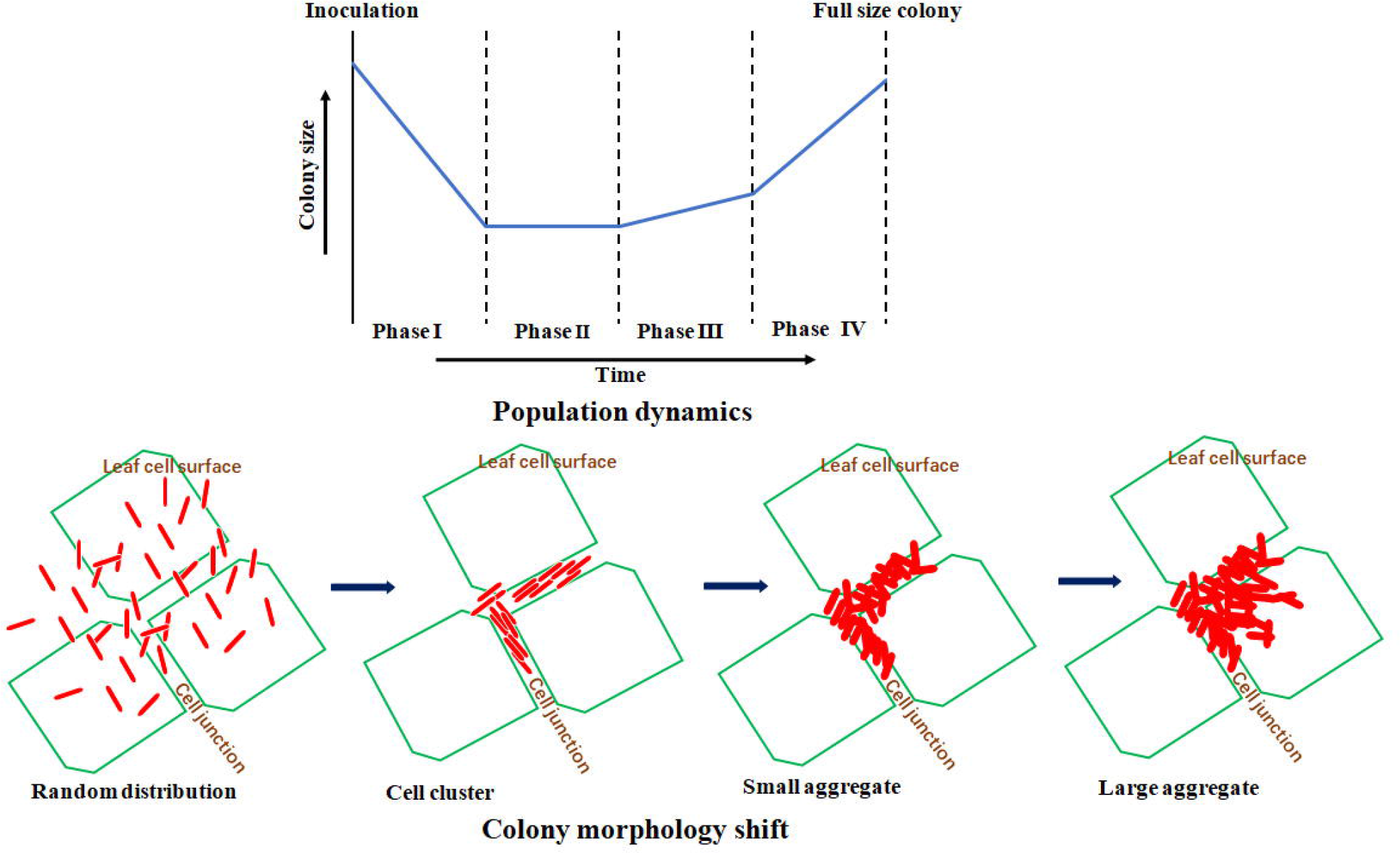
Colonization of *Rhodopseudomonas palustris* strain GJ-22 on leaf phyllosphere.

Phyllosphere bacteria form different sized aggregates or colonies at the junctions between plant epidermal cells or at the junctions along the veins or at the base of leaf trichrome (Beattie & Lindow, 1999). In this study, we observed that the preferential colonization location of strain GJ-22 was at the junctions between plant epidermal cells. In the early growth stages, most bacterial cells on leaf phyllosphere showed desiccation and then died. We speculate that the environment at the grooves formed by plant leaf epidermal cells provides bacterial cells a shelter with sufficient nutrients and water to survive and later to grow. Since the spaces provided by these grooves for bacteria growth are limited, most bacterial cells applied to plant canopy quickly desiccated and died. The survived cells started to multiple very slowly at the early colonization stage followed by a moderate multiplication to form Type 3 colonies. After entering the Phase III stage, the bacterial aggregates started to form its own microenvironments to capture nutrients from plant exudates, to maintain water within microenvironments, and to protect bacterial cells from lytic enzymes and/or toxic substances produced by plant or other microbes. This strategy is commonly used by phyllosphere epiphytes (Bulgarelli et al., 2013; Remus-Emsermann & Schlechter, 2018; Wilson et al., 1999) and can be used to explain the increased stress tolerance of strain GJ-22 after the Phase III stage.

Exopolysaccharides (EPSs) are the major ingredients mediating the adhesion between bacterial cells on plant surfaces to form bacterial aggregates (Limoli et al., 2015). EPSs can also provide protection to bacterial cells from various stresses. Production of EPSs was reported to be controlled via a cell-density-dependent manner known as quorum sensing (QS) in many Gram-negative bacteria (Marketon et al., 2003; Tan et al., 2014). Quorum sensing is a process used by bacteria to monitor their local population density and to regulate gene expressions at different population densities. Many Gram-negative bacteria also use QS to regulate their colony morphology that are vital for successful bacterial colonization and habitat modification on plants (Quiñones et al., 2005; Quiñones et al., 2004). It was suggested that when bacterial cell density at the Phase II stage reached a certain threshold, the QS system started to up-regulate EPS production to facilitate aggregate formation. *R. palustris* was reported to possess a *rpaI-rpaR* gene pair-based QS system (Schaefer et al., 2008). The rpaI protein utilizes a plant produced *p*-coumarate to synthesize a *p*-coumaroyl-homoserine lactone (*p*C-HSL) signal (Hidetada et al., 2012; Lindemann et al., 2011). To date, how *rpaI-rpaR* gene pair-based QS system regulates EPS production in *R. palustris* is unknown. The availability of nutrients on plant phyllosphere is likely the major determinant of epiphytic colonization. Numerous evidence have indicated that plant epiphytes can modify their habitats through modification of nutrient concentrations on plant surfaces (Beattie & Lindow, 1999). The size expansion of strain GJ-22 colonies at the Phase IV stage suggested a better nutrient condition during this phase. Strain GJ-22 was reported to produce IAA as a signal to promote plant growth (Su et al., 2017)(Pin et al., 2017). This phytohormone was also reported to induce release of nutrients, including saccharides, from plant cells (Limtong & Koowadjanakul, 2012; Spaepen et al., 2007). We speculate that the formation of bacterial aggregates may up-regulate IAA production in *R. palustris* and later infiltrate into plant cells.

In conclusion, the results obtained from this study revealed the process of *R. palustris* strain GJ-22 colony establishment. Because bacterial stress tolerance, the colony size expansion, and priming of ISR in plant to TMV infection all occurred after the formation of large sized bacterial colonies, it is likely that formation of bacterial aggregate has an important role on GJ-22 epiphytic fitness. Further investigations are needed to elucidate the mechanism(s) controlling aggregate formation. Our results can benefit the design of a better application method to deploy this bacteria to plant phyllosphere.

## Acknowledgements

This research was supported by the National Key R&D Program of China (2017YFD0200400), the National Science Foundation of China (31701764), Hunan agricultural science and technology innovation fund (2017GC04), Hunan Natural Science Foundation (2017JJ3169), and the Hunan Provincial key research and Development Plan (2016NK2199). The grant agencies had no role in study design, data collection and analysis, the decision to publish, or the preparation of this manuscript.

## Author contributions

PS, ZZ, DZ, XT, JC and LZ designed the experiments and analysed the data. PS, AC, WT, JP, and ZZ performed the experiments. XZ, XD, YL, MRH contributed reagents/materials. PS and DZ wrote the paper. The authors want to thank Dr. Xin Shun Ding (The Noble Foundation, Ardmore, USA [retired], for his help during preparation of this manuscript).

## Additional information

Competing financial interests: The authors declare no competing financial interests.

## Reference

Beattie, G.A., Lindow, S.E. 1999. Bacterial colonization of leaves: a spectrum of strategies. Phytopathology, 89(5), 353–359.

Berg, G. 2009. Plant-microbe interactions promoting plant growth and health: Perspectives for controlled use of microorganisms in agriculture. Applied Microbiology & Biotechnology, 84(1), 11–18.

Bonaterra, A., Cabrefiga, J., Camps, J., Montesinos, E. 2007. Increasing survival and efficacy of a bacterial biocontrol agent of fire blight of rosaceous plants by means of osmoadaptation. FEMS microbiology ecology, 61(1), 185–195.

Bulgarelli, D., Schlaeppi, K., Spaepen, S., van Themaat, E.V.L., Schulze-Lefert, P. 2013. Structure and functions of the bacterial microbiota of plants. Annual review of plant biology, 64, 807–838.

Duke, K.A., Becker, M.G., Girard, I.J., Millar, J.L., Fernando, W.D., Belmonte, M.F., Kievit, T.R. 2017. The biocontrol agent Pseudomonas chlororaphis PA23 primes Brassica napus defenses through distinct gene networks. BMC genomics, 18(1), 467.

Fawzy, Z., El-Magd, M.A., Li, Y., Ouyang, Z., Hoda, A. 2012. Influence of foliar application by EM “effective microorganisms”, amino acids and yeast on growth, yield and quality of two cultivars of onion plants under newly reclaimed soil. Journal of Agricultural Science (Toronto), 4(11), 26–39.

Haefele, D.M., Lindow, S.E. 1987. Flagellar motility confers epiphytic fitness advantages upon Pseudomonas syringae. Appl. Environ. Microbiol., 53(10), 2528–2533.

Hassani, M.A., Durán, P., Hacquard, S. 2018. Microbial interactions within the plant holobiont. Microbiome, 6(1), 58.

Hidetada, H., Schaefer, A.L., E Peter, G., Harwood, C.S. 2012. Anaerobic p-coumarate degradation by Rhodopseudomonas palustris and identification of CouR, a MarR repressor protein that binds p-coumaroyl coenzyme A. Journal of bacteriology, 194(8), 1960–7.

Idi, A., Nor, M.H.M., Wahab, M.F.A., Ibrahim, Z. 2015. Photosynthetic bacteria: an eco-friendly and cheap tool for bioremediation. Reviews in Environmental Science and Bio/Technology, 14(2), 271–285.

Kembel, S.W., O’Connor, T.K., Arnold, H.K., Hubbell, S.P., Wright, S.J., Green, J.L. 2014. Relationships between phyllosphere bacterial communities and plant functional traits in a neotropical forest. Proceedings of the National Academy of Sciences, 111(38), 13715–13720.

Kim, Y.C., Leveau, J., Gardener, B.B.M., Pierson, E.A., Pierson, L.S., Ryu, C.-M. 2011. The multifactorial basis for plant health promotion by plant-associated bacteria. Appl. Environ. Microbiol., 77(5), 1548–1555.

Lambais, M.R., Lucheta, A.R., Crowley, D.E. 2014. Bacterial community assemblages associated with the phyllosphere, dermosphere, and rhizosphere of tree species of the Atlantic forest are host taxon dependent. Microbial ecology, 68(3), 567–574.

Larimer, F.W., Chain, P., Hauser, L., Lamerdin, J., Malfatti, S., Do, L., Land, M.L., Pelletier, D.A., Beatty, J.T., Lang, A.S. 2004. Complete genome sequence of the metabolically versatile photosynthetic bacterium Rhodopseudomonas palustris. Nature biotechnology, 22(1), 55.

Limoli, D.H., Jones, C.J., Wozniak, D.J. 2015. Bacterial extracellular polysaccharides in biofilm formation and function. Microbiology spectrum, 3(3).

Limtong, S., Koowadjanakul, N. 2012. Yeasts from phylloplane and their capability to produce indole-3-acetic acid. World Journal of Microbiology and Biotechnology, 28(12), 3323–3335.

Lindemann, A., Pessi, G., Schaefer, A.L., Mattmann, M.E., Christensen, Q.H., Kessler, A., Hennecke, H., Blackwell, H.E., Greenberg, E.P., Harwood, C.S. 2011. Isovaleryl-homoserine lactone, an unusual branched-chain quorum-sensing signal from the soybean symbiont Bradyrhizobium japonicum. Proceedings of the National Academy of Sciences, 108(40), 16765–16770.

Lindow, S.E., Brandl, M.T. 2003. Microbiology of the phyllosphere. Appl. Environ. Microbiol., 69(4), 1875–1883.

Müller, D.B., Vogel, C., Bai, Y., Vorholt, J.A. 2016. The plant microbiota: systems-level insights and perspectives. Annual review of genetics, 50, 211–234.

Marketon, M.M., Glenn, S.A., Eberhard, A., González, J.E. 2003. Quorum sensing controls exopolysaccharide production in Sinorhizobium meliloti. Journal of bacteriology, 185(1), 325–331.

Natacha, B., Miriam, B.M., Martin, A., Vorholt, J.A. 2014. A synthetic community approach reveals plant genotypes affecting the phyllosphere microbiota. Plos Genetics, 10(4), e1004283.

Okinaka, Y., Yang, C.-H., Perna, N.T., Keen, N.T. 2002. Microarray profiling of Erwinia chrysanthemi 3937 genes that are regulated during plant infection. Molecular plant-microbe interactions, 15(7), 619–629.

Poonguzhali, S., Madhaiyan, M., Yim, W.-J., Kim, K.-A., Sa, T.-M. 2008. Colonization pattern of plant root and leaf surfaces visualized by use of green-fluorescent-marked strain of Methylobacterium suomiense and its persistence in rhizosphere. Applied microbiology and biotechnology, 78(6), 1033–1043.

Quiñones, B., Dulla, G., Lindow, S.E. 2005. Quorum sensing regulates exopolysaccharide production, motility, and virulence in Pseudomonas syringae. Molecular plant-microbe interactions, 18(7), 682–693.

Quiñones, B., Pujol, C.J., Lindow, S.E. 2004. Regulation of AHL production and its contribution to epiphytic fitness in Pseudomonas syringae. Molecular plant-microbe interactions, 17(5), 521–531.

Remus-Emsermann, M.N., Schlechter, R.O. 2018. Phyllosphere microbiology: at the interface between microbial individuals and the plant host. New Phytologist, 218(4), 1327–1333.

Sasaki, K., Watanabe, M., Suda, Y., Ishizuka, A., Noparatnaraporn, N. 2005. Applications of photosynthetic bacteria for medical fields. Journal of bioscience and bioengineering, 100(5), 481–488.

Schaefer, A.L., Greenberg, E., Oliver, C.M., Oda, Y., Huang, J.J., Bittan-Banin, G., Peres, C.M., Schmidt, S., Juhaszova, K., Sufrin, J.R. 2008. A new class of homoserine lactone quorum-sensing signals. Nature, 454(7204), 595.

Spaepen, S., Vanderleyden, J., Remans, R. 2007. Indole-3-acetic acid in microbial and microorganism-plant signaling. FEMS microbiology reviews, 31(4), 425–448.

Su, P., Tan, X., Li, C., Zhang, D., Cheng, J.e., Zhang, S., Zhou, X., Yan, Q., Peng, J., Zhang, Z. 2017. Photosynthetic bacterium R hodopseudomonas palustris GJ-22 induces systemic resistance against viruses. Microbial biotechnology, 10(3), 612–624.

Tan, C.H., Koh, K.S., Xie, C., Tay, M., Zhou, Y., Williams, R., Ng, W.J., Rice, S.A., Kjelleberg, S. 2014. The role of quorum sensing signalling in EPS production and the assembly of a sludge community into aerobic granules. The ISME journal, 8(6), 1186.

Vorholt, J.A. 2012. Microbial life in the phyllosphere. Nature Reviews Microbiology, 10(12), 828–840.

Wilson, M., Hirano, S., Lindow, S. 1999. Location and survival of leaf-associated bacteria in relation to pathogenicity and potential for growth within the leaf. Appl. Environ. Microbiol., 65(4), 1435–1443.

Wong, W.-T., Tseng, C.-H., Hsu, S.-H., Lur, H.-S., Mo, C.-W., Huang, C.-N., Hsu, S.-C., Lee, K.-T., Liu, C.-T. 2014. Promoting effects of a single Rhodopseudomonas palustris inoculant on plant growth by Brassica rapa chinensis under low fertilizer input. Microbes and environments, ME14056.

Wu, J., Wang, Y., Lin, X. 2013. Purple phototrophic bacterium enhances stevioside yield by Stevia rebaudiana Bertoni via foliar spray and rhizosphere irrigation. PloS one, 8(6), e67644.

